# Cellular management of Zinc in group B *Streptococcus* supports bacterial resistance against metal intoxication and promotes disseminated infection

**DOI:** 10.1101/2021.02.04.429868

**Authors:** Matthew J. Sullivan, Kelvin G. K. Goh, Glen C. Ulett

## Abstract

Zinc (Zn) is an essential trace element for normal bacterial physiology but divergently, can intoxicate bacteria at high concentrations. Here, we define the molecular systems for Zn detoxification in *Streptococcus agalactiae*, also known as group B Streptococcus, and examine the effects of resistance to Zn stress on virulence. We compared the growth of wild-type bacteria and mutants deleted for the Zn exporter, *czcD*, and the response regulator, *sczA*, using Zn-stress conditions *in vitro.* Macrophage antibiotic protection assays and a mouse model of disseminated infection were used to assess virulence. Global bacterial transcriptional responses to Zn stress were defined by RNA-sequencing and qRTPCR. *czcD* and *sczA* enabled *S. agalactiae* to survive Zn stress, with the putative CzcD efflux system activated by SczA. Additional genes activated in response to Zn stress encompassed divalent cation transporters that contribute to regulation of Mn and Fe homeostasis. *In vivo*, the *czcD-sczA* Zn-management axis supported virulence in the blood, heart, liver and bladder. Additionally, several genes not previously linked to Zn stress in any bacterium, including most notably, *arcA* for arginine deamination also mediated resistance to Zn stress; representing a novel molecular mechanism of bacterial resistance to metal intoxication. Taken together, these findings show that *S. agalactiae* responds to Zn stress by *sczA* regulation of *czcD*, with additional novel mechanisms of resistance supported by *arcA*, encoding arginine deaminase. Cellular management of Zn stress in *S. agalactiae* supports virulence by facilitating bacterial survival in the host during systemic infection.

**Importance Statement:** *Streptococcus agalactiae*, also known as group B streptococcus, is an opportunistic pathogen that causes various diseases in humans and animals. This bacterium has genetic systems that enable Zinc (Zn) detoxification in environments of metal stress, but these systems remain largely undefined. Using a combination of genomic, genetic and cellular assays we show that this pathogen controls Zn export through CzcD to manage Zn stress, and utilizes a system of arginine deamination never previously linked to metal stress responses in bacteria to survive metal intoxication. We show that these systems are crucial for survival of *S. agalactiae in vitro* during Zn stress and also enhance virulence during systemic infection in mice. These discoveries establish new molecular mechanisms of resistance to metal intoxication in bacteria; we suggest these mechanisms are likely to operate in other bacteria as a way to sustain microbial survival in conditions of metal stress, including in host environments.

## Introduction

Inside living cells, zinc (Zn) is an essential cofactor for metalloenzymes (1, 2) but is toxic at high concentrations, as can be encountered by bacteria inside phagocytes (3, 4). The double-edged sword of essentiality and toxicity of Zn to bacteria is a burgeoning area of research due to potential for antimicrobial applications (5–7). In the host, bacterial pathogens employ distinct mechanisms for internalizing essential Zn (8, 9), and in turn, the host can restrict Zn availability as an antimicrobial strategy (10). Phagocytes can mobilise cellular Zn to expose internalized bacteria to metal concentrations that are antimicrobial (5, 11, 12). Host-driven Zn intoxication (defined as an excess of extracellular Zn) of bacterial pathogens can involve ablation of uptake of essential Mn (13), a compromised bacterial response to oxidative stress (14), or disrupted central carbon metabolism (15). Some bacteria can evade metal intoxication by mechanisms that involve metal efflux (16).

In streptococci, a specific genetic system manages Zn homeostasis by regulating metal import and export (13, 17). In Pneumococcus and *Streptococcus pyogenes*, systems for Zn efflux pair a Zn-sensing transcriptional response regulator *(sczA)* with a Zn efflux transporter, encoded by *czcD* (17, 18). Here, we studied Zn management in *Streptococcus agalactiae*, also known as group B *Streptococcus*, which is an important opportunistic pathogen that has undefined Zn detoxification systems and is associated with distinct disease aetiologies compared to other streptococci. We establish a role for CzcD, as well as other novel factors, in mediating *S. agalactiae* resistance to Zn stress and virulence.

## Materials and Methods

### Bacterial strains, plasmids and growth conditions

*S. agalactiae, E. coli* and plasmids used are listed in Supplementary Table 1. *S. agalactiae* was routinely grown in Todd-Hewitt Broth (THB) or on TH agar (1.5% w/v). *E. coli* was grown in Lysogeny Broth (LB) or on LB agar. Media were supplemented with antibiotics (spectinomycin (Sp) 100μg/mL; chloramphenicol (Cm) 10 μg/mL), as indicated. Growth assays used 200μL culture volumes in 96- well plates (Greneir) sealed using Breathe-Easy^®^ membranes (Sigma Aldrich) and measured attenuance (*D*, at 600nm) using a ClarioSTAR multimode plate reader (BMG Labtech). Media for growth assays were THB and a modified Chemically-Defined Medium (CDM) (9) (with 1g/L glucose, 0.11g/L pyruvate and 50μg/L L-cysteine), supplemented with Zn (supplied as ZnSO_4_) as indicated. For attenuance baseline correction, control wells without bacteria were included for Zn in media alone.

### DNA extraction and genetic modification of *S. agalactiae*

Plasmid DNA was isolated using miniprep kits (QIAGEN), with modifications for *S. agalactiae* as described elsewhere (19). Deletion of *czcD* (CHF17_00567 / CHF17_RS02855 = ASZ00854.1) was constructed by allelic exchange with a Cm cassette using pHY304aad9 as described previously (20). Deletions in *sczA* and *arcA* were generated similarly but without Cm-markers. Constructs, complement plasmids and primers are listed in Supplementary Table 2. Constructs for *sczA* and *arcA* deletions were made with DNA synthesised in pUC57 by Genscript (USA) prior to subcloning into pHY304aad9. Mutants were validated by PCR using primers external to the mutation site and DNA sequencing.

### Expression system for mCherry in *S. agalactiae*

Plasmid pGU2665 was designed for expressing mCherry *in trans* in *S. agalactiae* from the pCP25 promoter (21) cloned into pDL278 (22) (Supplementary Figure 1). Plasmid DNA was manipulated and ligation reactions were performed essentially as described elsewhere (19). We verified pGU2665 by restriction analysis and sequencing using primers listed in Supplementary Table 2. Sequence reads were mapped and assembled using Sequencher software. Electroporation of *S. agalactiae* and selection of transformants was performed as described previously (20). Plasmid stability assays were performed as described previously (19).

### RNA extraction, qRTPCR

For Zn exposure experiments, 1mL overnight THB cultures were back-diluted 1/100 in 100mL of THB (prewarmed at 37°C in 250mL Erlenmeyer flasks) supplemented with 0.25, 0.5, 1.0 or 1.5mM Zn. Cultures were grown shaking (200rpm) at 37°C; after exactly 2.5h, 10-50mL volumes containing approximately 500 million mid-log bacteria were harvested; RNA was preserved and isolated as described previously (23). RNA quality was analysed by RNA LabChip using GX Touch (Perkin Elmer). RNA (1000ng) was reverse-transcribed using Superscript IV according to manufacturer’s instructions (Life Technologies) and cDNA was diluted 1:50 in water prior to qPCR. Primers (Supplementary Table 2) were designed using Primer3 Plus (24, 25) to quantify transcripts using Universal SYBR Green Supermix (Bio-Rad) using a Quantstudio 6 Flex (Applied Biosystems) system in accordance with MIQE guidelines (26). Standard curves were generated using five point serial dilutions of genomic DNA (5-fold) from WT *S. agalactiae* 874391 (27). Expression ratios were calculated using C_T_ values and primer efficiencies as described elsewhere (28) using *dnaN*, encoding DNA polymerase III β-subunit as housekeeper.

### Whole bacterial cell metal content determination

Metal content in cells was determined as described (14) with minor modifications. Cultures were prepared essentially as described for *RNA extraction, qRTPCR* with the following modifications; THB medium was supplemented with 0.25 mM ZnSO_4_ or not supplemented (Ctrl), and following exposure for 2.5h, bacteria were harvested by centrifugation at 4122 x g at 4°C. Cell pellets were washed 3 times in PBS + 5mM EDTA to remove extracellular metals, followed by 3 washes in PBS. Pelleted cells were dried overnight at 80°C and resuspended in 1mL of 32.5% nitric acid and incubated at 95°C for 1h. The metal ion containing supernatant was collected by centrifugation (14,000 x g, 30min) and diluted to a final concentration of 3.25% nitric acid for metal content determination using inductively coupled plasma optical emission spectroscopy (ICP-OES). ICP-OES was carried out on an Agilent 720 ICP-OES with axial torch, OneNeb concentric nebulizer and Agilent single pass glass cyclone spray chamber. The power was 1.4kW with 0.75L/min nebulizer gas, 15L/min plasma gas and 1.5L/min auxiliary gas flow. Zn was analyzed at 213.85nm, Cu at 324.75nm, Fe at 259.94nm, Mn at 257.61nm with detection limits at <1.1ppm. The final quantity of each metal was normalised using dry weight biomass of the cell pellet prior to nitric acid digestion, expressed as μg.g ^-1^dry weight. Baseline concentrations were determined to be 10.9 ± 0.07μM Zn in THB medium, and 0.11 ± 0.03μM Zn in CDM medium from at least three independent assays.

### RNA sequencing and bioinformatics

Cultures were prepared as described above for *RNA extraction, qRTPCR.* RNase-free DNase-treated RNA that passed Bioanalyzer 2100 (Agilent) analysis was used for RNA sequencing (RNA-seq) using the Illumina NextSeq 500 platform. We used TruSeq library generation kits (Illumina, San Diego, California). Library construction consisted of random fragmentation of the poly(A) mRNA, followed by cDNA production using random primers. The ends of the cDNA were repaired and A-tailed, and adaptors were ligated for indexing (with up to 12 different barcodes per lane) during the sequencing runs. The cDNA libraries were quantitated using qPCR in a Roche LightCycler 480 with the Kapa Biosystems kit (Kapa Biosystems, Woburn, Massachusetts) prior to cluster generation. Clusters were generated to yield approximately 725K–825K clusters/mm^2^. Cluster density and quality was determined during the run after the first base addition parameters were assessed. We ran paired-end 2 × 75–bp sequencing runs to align the cDNA sequences to the reference genome. For data preprocessing and bioinformatics, STAR (version 2.7.3a) was used to align the raw RNA sequencing fastq reads to the WT *S. agalactiae* 874391 reference genome (27). HTSeq-count, version 0.11.1, was used to estimate transcript abundances (29). DESeq2 was then used to normalized and test for differential expression and regulation. Genes that met certain criteria (i.e. fold change of > ±2.0, q value of <0.05) were accepted as significantly altered (30). Raw and processed data were deposited in Gene Expression Omnibus (accession no. GSE161127).

### Mammalian cell culture

J774A.1 murine macrophages or U937 human monocyte-derived macrophages (MDMs) were grown in RPMI and seeded (10^5^) into the wells of a 96-well tissue culture-treated plate (Falcon) essentially as described elsewhere (31, 32), except that U937 MDMs were differentiated by exposure to 30ng/mL phorbol 12- myristate 13-acetate (PMA) for 48h and cells subsequently rested in media without PMA for 72h to enhance morphological and phenotypic markers of MDMs (33). A multiplicity of infection (MOI) of 100 bacteria:macrophage for 1h was used in RPMI without antibiotics. Non-adherent bacteria were removed by five washes of 200μL PBS using a Well Wash Versa (Thermo Scientific). RPMI containing 250U/mL penicillin, streptomycin (Gibco) and 50μg/mL gentamicin (Sigma-Aldrich) were used for antibiotic protection assays to quantify intracellular bacteria as described previously (32). At 1h, 24h or 48h after infection, monolayers were washed five times with 200μL PBS and lysed by brief exposure to 50μL of 2% trypsin and 0.02% Triton-X-100 (10min) prior to dilution with 150μL PBS and estimation of CFU/mL by serial dilution and plate counts on agar.

### Fluorescence microscopy

Fifty thousand J774A.1 cells were seeded into 8-well LabTek II chamber slides (Nunc) and infected with mCherry-*S. agalactiae* for 24h. Non-adherent bacteria were removed by three washes of 200μL PBS and fixed for 15min at 37°C using 4% paraformaldehyde (w/vol). Monolayers were stained for Zn^2+^ using 5μM FluoZin-3^TM^ AM (Life Technologies) for 30min at 37°C, and subsequently, for DNA using Hoechst 33258 for 5min. Fixed stained cells were washed twice in PBS and mounted using n-propyl gallate (n-pg) mounting medium (0.2% n-pg in 9:1 glycerol/PBS). mCherry-*S. agalactiae* was visualized using a Zeiss AxioImager.M2 microscope (Carl Zeiss MicroImaging) fitted with PlanApochromat X20/0.8 and X63/1.40 objective lenses and an AxioCam MRm Rev.3 camera. Images of cells were captured with 63HE, 44 and 49 filter sets (to detect mCherry (587nm, 610nm), FluoZin^TM^-3 (494nm, 518nm) and Hoechst 33258 (352nm, 461nm) fluorescence, respectively, with excitation and emission spectra listed consecutively for each) and Zen Pro (version 2) software.

### Animals and Ethics statement

Virulence was tested using a mouse model of disseminated infection based on intravenous challenge with 10^7^ *S. agalactiae* as described elsewhere (34). This study was carried out in accordance with the guidelines of the Australian National Health and Medical Research Council. The Griffith University Animal Ethics Committee reviewed and approved all experimental protocols for animal usage according to the guidelines of the National Health and Medical Research Council (approval: MSC/01/18/AEC).

### Statistical methods

All statistical analyses used GraphPad Prism V8 and are defined in respective Figure Legends. Statistical significance was accepted at P values of ≤0.05.

## Results

### Excess Zn impedes *S. agalactiae* growth and disturbs cell physiology

Initial assays analyzed the growth of wild-type (WT) *S. agalactiae* strain 874391 (ST-17; Serotype III) in a nutrient-rich Todd-Hewitt broth (THB) supplemented with moderate (0.5 mM), high (1.0 mM) and excess (1.5 mM) levels of Zn. High and excess Zn (≥1 mM) delayed exponential growth of the WT with significant attenuation of the growth rate and final biomass yield (Figure 1A). An isogenic *ΔczcD* mutant strain was significantly more susceptible to Zn ≥ 1mM (Figure 1A). Full-length *czcD* supplied *in trans* to the mutant (Δ*czcD*::*czcD*) restored growth to WT levels in Zn stress (Figure 1A). Spot test assays of the bacteria on agar containing 0, 0.5, 1.0 and 1.5 mM Zn showed similar levels of susceptibility of the WT and *ΔczcD* strains (Figure 1B) on solid medium (Figure 1B) compared to planktonic growth.

**Figure 1.**
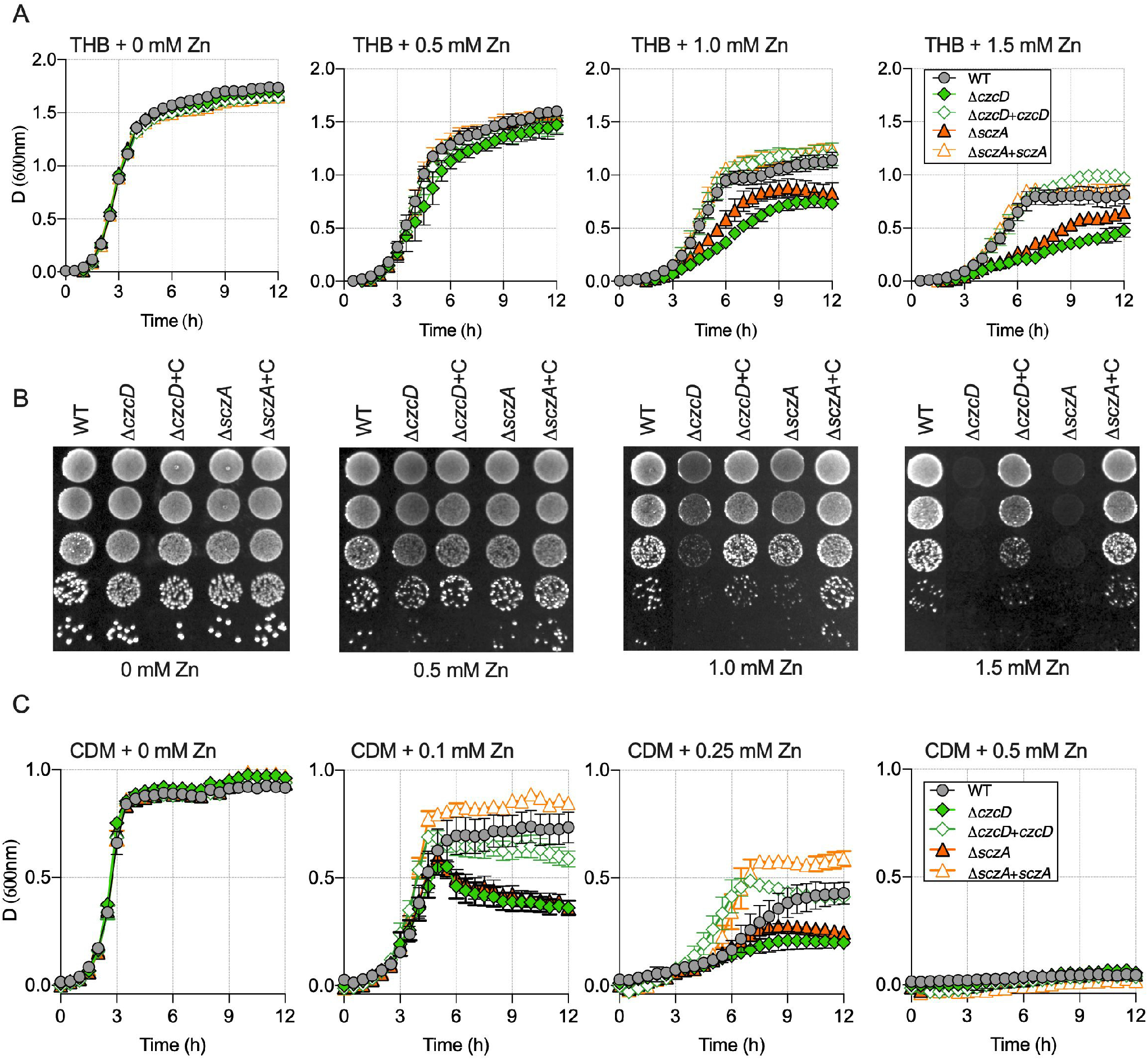
Comparison of the growth WT *S. agalactiae, ΔczcD* and *ΔsczA* isogenic mutants and complemented strains. A, Growth curves of 874391 and mutant strains as indicated, in Todd-Hewitt broth (THB) supplemented with 0, 0.5, 1.0 and 1.5 mM Zn. B, Analysis of growth of *S. agalactiae* strains on solid TH medium with increasing Zn concentrations by serial dilution and droplets on agar. Composite images are merged from multiple agar plates. C, Growth curves of 874391 and mutant strains as indicated, in Chemically Defined Medium (CDM) supplemented with 0, 0.1, 0.25 and 0.5 mM Zn. Data shown are mean measurements of attenuance (*D*; at 600nm) from 3-4 independent experiments, bars show S.E.M. Attenuance measurements used well-scan mode with a 3mm 5×5 scan matrix, 5 flashes per point and path length correction of 5.88mm, 300rpm agitation, every 30min

We next used a nutrient limited medium to examine Zn stress in *S. agalactiae* in a modified Chemically Defined minimal Medium (CDM), which is likely to more closely reflect a host-environment (nutrient-limited). CDM contained low basal levels of Zn (0.11 ± 0.03 μM), as determined by ICP-OES. Growth assays of *S. agalactiae* in CDM ± Zn supplementation revealed markedly enhanced Zn-induced toxicity compared to THB, with Zn totally bacteriostatic to all strains at 0.5 mM (Figure 1C). Similar to THB, growth of the *ΔczcD* mutant was significantly inhibited compared to the WT upon exposure to ≥ 0.1 mM in CDM (Figure 1C).

### Regulation of Zn-efflux in *S. agalactiae*

The capacity of Zn stress to induce expression of *czcD* for Zn export was examined by analyzing mid-log phase *S. agalactiae* exposed to 0.25-1.5 mM Zn for 2.5h prior to qRTPCR quantification of *czcD. czcD* was significantly upregulated in response to Zn (3-fold – 18.9-fold) in a manner that was titratable with the Zn concentration (Figure 2A); and consistent with a role for CzcD in responding to extracellular Zn. Interestingly, we identified a candidate gene, divergent from *czcD*, that encodes a putative Zn-responsive activator in the TetR-family of transcription factors, termed **<u>s</u>**treptococcal ***<u>cz</u>***cD **<u>A</u>**ctivator, or *szcA* (35). Similar to *ΔczcD S. agalactiae*, a mutant deficient in *sczA* (Δ*sczA*) was markedly more susceptibility to Zn stress compared to WT in both nutrient-rich and -limited conditions; the attenuation was restored by complementation *in trans* (Figure 1).

**Figure 2.**
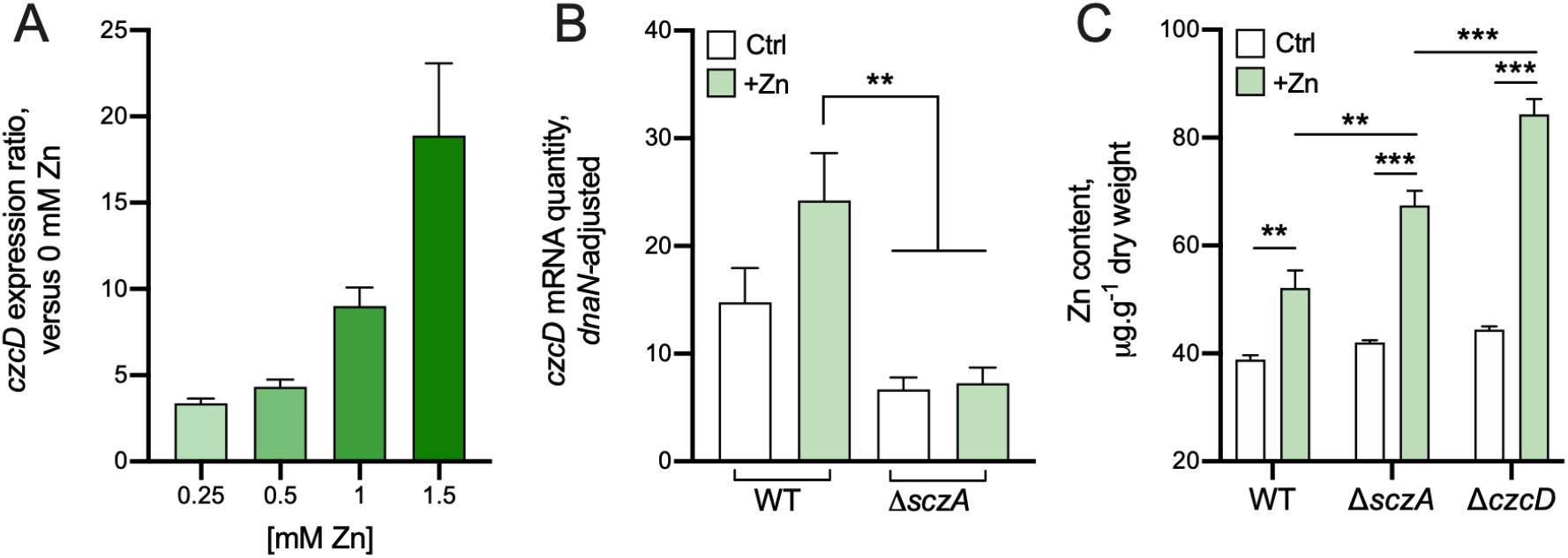
Expression analysis of *czcD* and intracellular Zn content in *S agalactiae* strains. A, Expression ratio of *czcD* quantified by qRTPCR in THB medium containing 0.25, 0.5, 1.0 and 1.5 mM Zn, compared to THB without Zn. B, Relative *czcD* transcripts were quantified in WT and *ΔsczA* strains with and without Zn supplementation. C, Intracellular accumulation of Zn was compared with and without Zn supplementation in WT, *ΔsczA* and *ΔczcD* strains. Ratios in A were calculated as described previously (28) using C_T_ values, primer efficiencies and housekeeping *dnaN.* In B and C, Ctrl = THB + 0 mM; +Zn = THB + 0.25 mM Zn. Bars show means and S.E.M from 3-4 independent experiments and compared by One-way ANOVA with Holm-Sidak multiple comparisons (**P < 0.01; *** P <0.001).

To confirm the role of SczA as a Zn-responsive activator of *czcD* expression, we quantified *czcD* mRNA in *ΔsczA S. agalactiae* exposed to 0.25 mM Zn (sub-inhibitory; to enable comparisons independent of metabolic state, and alleviate potential bias from any discordant Zn stress between WT and mutants with varied resistance phenotypes). Deletion of *sczA* significantly perturbed activation of *czcD* in response to Zn (Figure 2B), consistent with previous reports of SczA functioning as a Zn-dependent activator of *czcD* in *S. pyogenes* and *S. pneumoniae* (17, 18).

### Intracellular Zn content in *S. agalactiae* during Zn stress

We analyzed accumulation of Zn ions within the bacteria following Zn stress by using growth conditions equivalent to those for transcriptional experiments; WT, *ΔczcD* and *ΔsczA S. agalactiae* were exposed to 0.25 mM Zn for 2.5 h prior to quantifying intracellular metal content by ICP-OES. In the WT strain, Zn stress caused accumulation of intracellular Zn (52.1 vs 38.8 μg Zn/g dry weight) compared to non-exposed control cultures. In addition, intracellular Zn contents in *ΔczcD* and *ΔsczA* mutants exposed to the same conditions were significantly enhanced (84.3 and 67.5 μg Zn/g dry weight, respectively) compared to WT in Zn stress or control incubations of the mutant strains without Zn (Figure 2C). These findings are consistent with roles for CzcD and SczA as mediators of Zn efflux.

### Resistance of intracellular *S. agalactiae* to Zn stress in macrophages

To determine if Zn efflux systems support survival of *S. agalactiae* in phagocytic cells, we used murine macrophages and human monocyte-derived macrophagelike cells in antibiotic protection assays at 1h, 24h and 48h of incubation. Viable intracellular *S. agalactiae* were reduced in number over the time course in human and murine cells, but no significant differences between WT, *ΔczcD* and *ΔsczA* strains were detected (Figure 3A). To examine host-mediated Zn release inside phagocytic cells, we developed a system for expressing mCherry in *S. agalactiae.* We used this in concert with a Zn-binding fluorophore (FluoZin^TM^-3 AM) to visualize intracellular *S. agalactiae* inside J774A.1 cells. Despite no effect of Zn efflux mutants on survival, we observed that *S. agalactiae* induced robust mobilization of Zn inside host cells at the 24h timepoint, compared to noninfected control incubations (Figure 3B) as shown by enhanced detection and distribution of free Zn using FluoZin^TM^-3AM.

**Figure 3.**
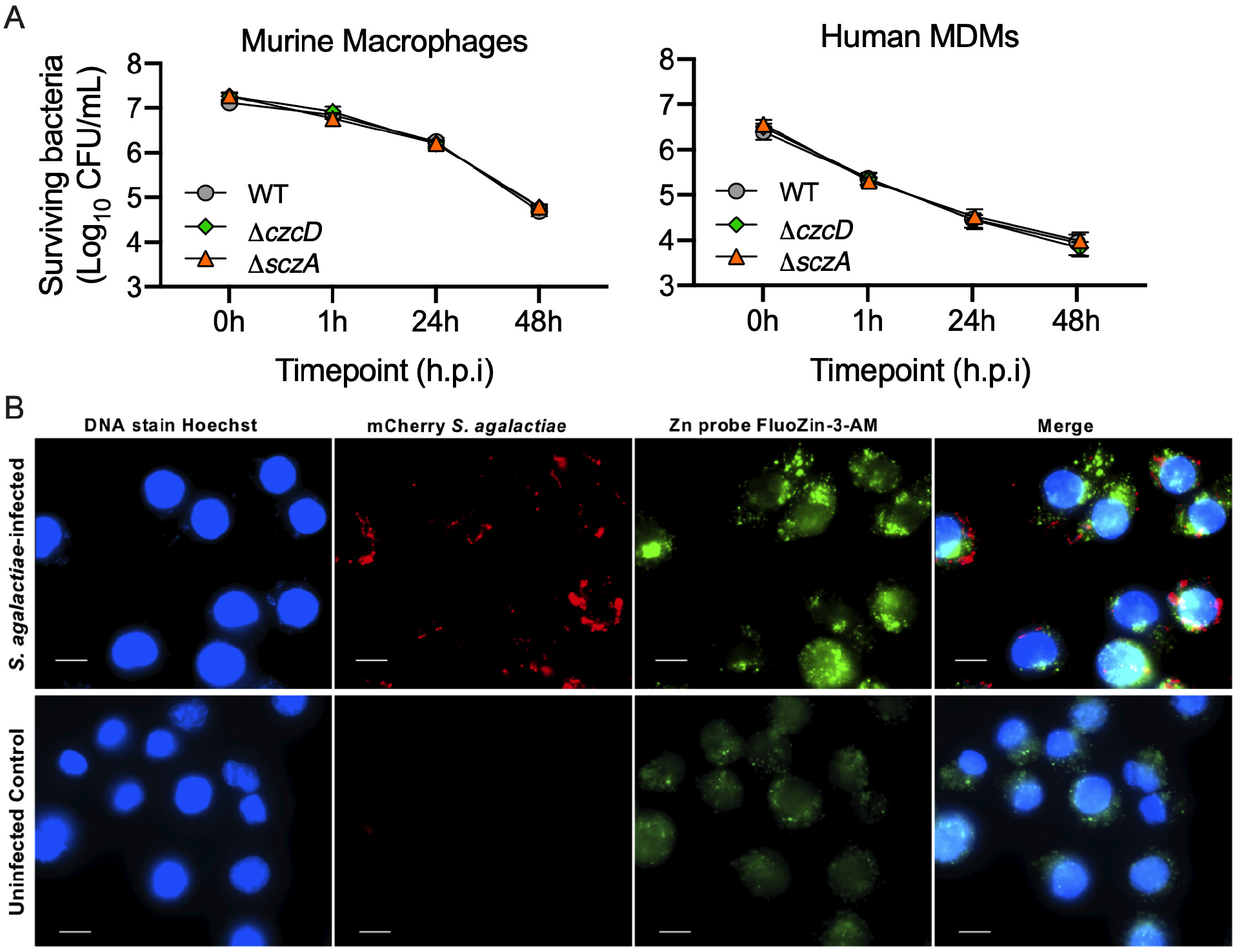
Interactions of *S. agalactiae* with macrophages and consequential mobilisation of Zn. A, Gentamicin protection assays with WT, *ΔsczA* and *ΔczcD* strains and mouse (J774A.1) or human (U937 monocyte-derived macrophage like) macrophages, with surviving bacteria quantified at 0, 1, 24 and 48 hours post infection (h.p.i). Data are means and S.E.M of 4-5 independent experiments. B, Fluorescence imaging of the release of free Zn (detected using FluoZin^TM^-3 AM) by J774A.1 cells infected with WT *S. agalactiae* expressing mCherry from pGU2665, compared to non-infected J774A.1 cells at 24 h post inoculation. DNA was stained using Hoechst 33258. Scale bars = 10μm.

### *S. agalactiae* Zn efflux systems contribute to virulence *in vivo*

To examine the contribution of Zn efflux to *in vivo* colonization of *S. agalactiae*, we used a murine model of disseminated infection to monitor tissue and bloodstream burdens (34). In mice that were challenged intravenously with 10^7^ bacteria and monitored for bacterial load, we detected significantly fewer *ΔczcD* mutant in the liver (P= 0.046) and bladder (P= 0.025), compared to the WT at 24 h post-inoculation (Figure 4A). No differences were observed between counts of the WT and *ΔczcD* mutant in the brain, blood, heart, lungs, kidneys, or spleen (data not shown). In addition, significantly fewer *ΔsczA* mutant were recovered from the blood (P= 0.039) and heart (P= 0.032) compared to the WT (Figure 4B), indicating a modest but statistically significant role for cellular management of Zn via *czcD* and *sczA* in supporting disseminated infection *in vivo.*

**Figure 4.**
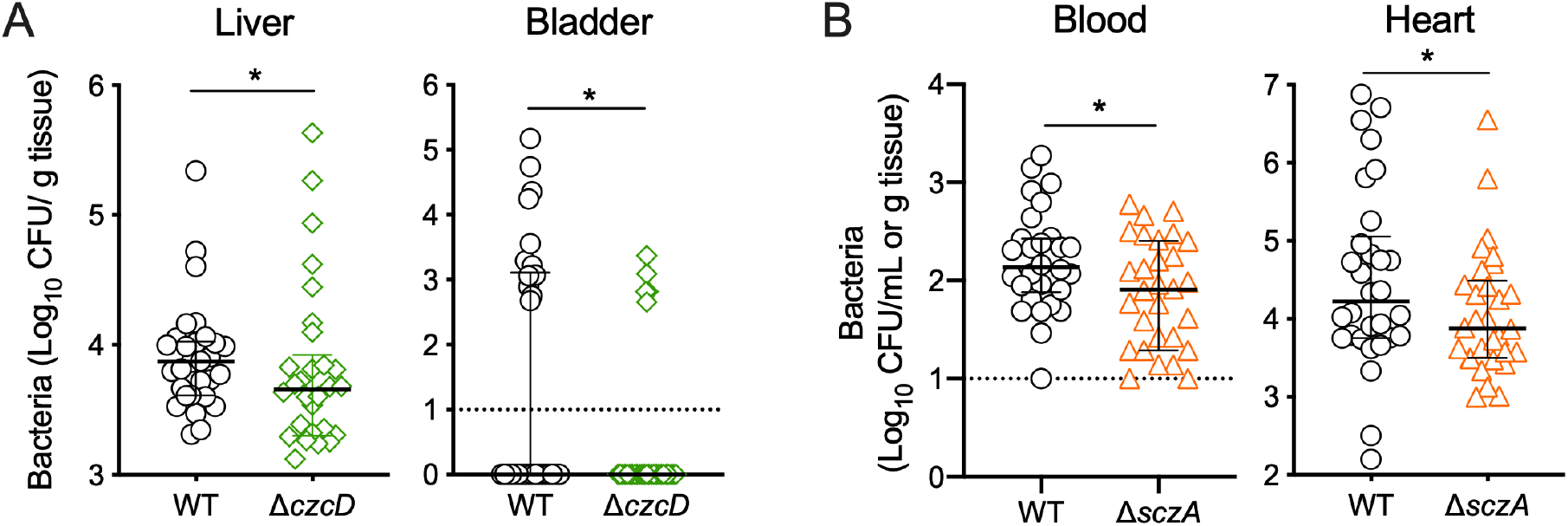
Virulence of WT (grey circles), *ΔczcD* (green diamonds) and *ΔsczA S. agalactiae* (orange triangles) in a mouse model of disseminated infection. C57BL/6 mice (6-8 weeks old) were intravenously injected with 10^7^ bacteria; bacteremia and disseminated spread to the heart (A), liver, and bladder (B) were monitored at 24h post infection. CFU were enumerated and counts were normalized using tissue mass in g. Lines and bars show median and interquartile ranges and data are pooled from 3 independent experiments each containing n=10 mice with mutant strains compared using Mann-Whitney U-tests to WT colonization data (*P < 0.05, **P < 0.01).

### *S. agalactiae*Zn stress transcriptome reveals new mediators of resistance

The transcriptome of *S. agalactiae* in response to Zn stress was used to define the global response of this organism to externally applied Zn. RNA-Seq identified 567 genes that were differentially expressed upon exposure to 0.25 mM Zn (229 up-, 238 down-regulated; ±2-fold, P-adj <0.05, n=4 biological replicates) in WT *S. agalactiae* (Figure 5A and Supplementary Table 3). In addition to up-regulation of *czcD*, we detected up-regulation of several putative nickel (Ni) and manganese (Mn) transport loci *(nikABCD, mntH2*, and *mtsABC*, respectively), and down-regulation of an iron (Fe) efflux system *(fetAB)* and Zn-importing *adcA* (Figure 5B). Interestingly, some these transport genes were recently implicated in *S. agalactiae* survival against host-derived calprotectin (36), which mediates Zn starvation, as opposed to Zn intoxication.

**Figure 5.**
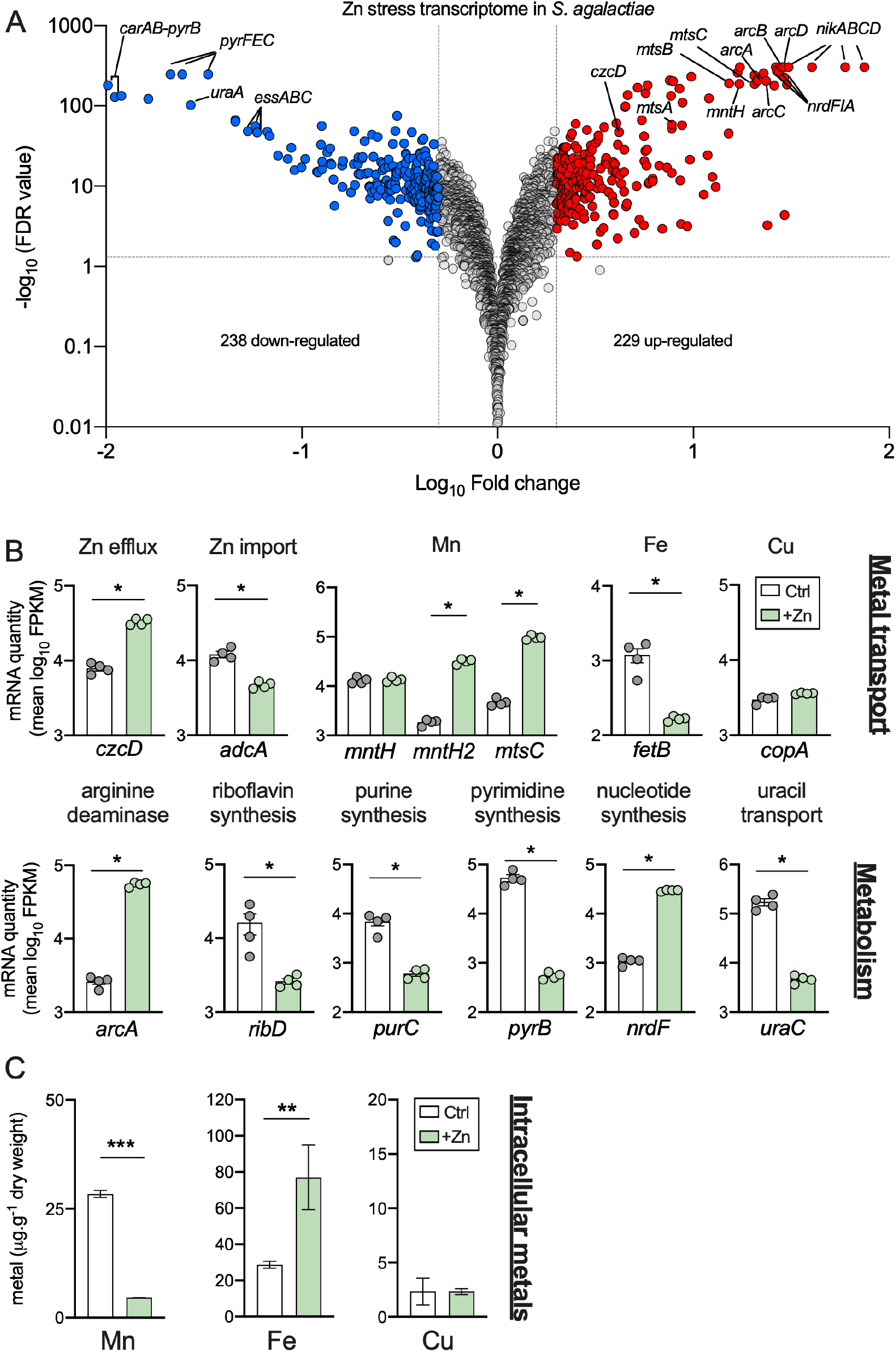
Transcriptomic analysis of *S. agalactiae* in Zn stress. A, Volcano plot showing data from RNASeq analysis of WT *S. agalactiae* cultures containing 0.25 mM Zn compared to non-exposed controls. Transcripts up- or down-regulated in response to Zn (*n*=4, >± 2-fold, FDR <0.05) are highlighted in red and blue, respectively. Dotted lines show False discovery rate (FDR; q-value) and fold-change cut-offs. Grey points indicate genes that were unchanged. Selected genes are identified individually with black lines. B, Expression of selected genes from RNASeq analyses showing mean <u>F</u>ragments <u>P</u>er <u>K</u>ilobase of transcript per Million mapped reads (FPKM) values for each condition, with predicted function as indicated. Data were compared with DESeq2 (* P-adj < 0.05 and ±2-fold). C, Intracellular accumulation of Mn, Fe and Cu were compared with and without Zn supplementation in WT *S. agalactiae* using ICP-OES. Bars show means and S.E.M from 3-4 independent experiments and compared by unpaired *t-*tests (**P < 0.01; *** P <0.001).

To provide functional insight into the observed changes in metal transporters, we analysed cellular metal content of *S. agalactiae* during Zn stress, for other second row transition metals, including Mn, Fe, Ni and Cu. We saw significant reduction in Mn levels, enhanced cellular Fe, no difference in Cu levels, and Ni was below the detection limit (0.8 ppm) (Figure 5C), when comparing cultures of WT bacteria under Zn stress (0.25 mM) to non-supplemented controls (THB media only).

In addition to metal transporters, we detected modulation of several metabolism-related gene clusters that have not previously been linked to Zn stress in bacteria, encompassing *de novo* nucleotide synthesis and import systems *(carAB, pyrB, pyrFEC, pur* genes, *uraA, nrdFIA, guaC)* and riboflavin synthesis *(ribDEAH)* loci (Figure 5 and Supplementary Table 3). Interestingly, a putative arginine deaminase system (ADI), encoded by the *arcABCD* genes, was significantly up-regulated (21-29-fold) under Zn intoxication *(arcA* highlighted in Figure 5B). The *arcABC* genes encode arginine deaminase (ArcA), ornithine carbamoyltransferase (ArcB) and carbamate kinase (ArcC), with *arcD* encoding an ornithine/arginine antiporter. This system functions to produce ammonia and ATP from the conversion of arginine to ornithine, as characterised in *S. pneumoniae* (37) and is responsive to numerous stimuli (38). We generated an isogenic *ΔarcA* mutant to examine a potential role for ADI in Zn stress resistance in *S. agalactiae.* Comparing the growth of the *ΔarcA* mutant to WT in nutrient-rich THB + 1 mM Zn and nutrient-limiting CDM + 0.1 mM Zn (conditions in which *ΔczcD/ΔsczA* strains were perturbed) revealed significant attenuation of the *ΔarcA* strain for growth under Zn stress (Figure 6), noting a modest reduction in growth of the *ΔarcA* strain vs WT in control conditions (without added Zn). Finally, *S. agalactiae* modulated several genes encoding classical virulence and/or immunogenic factors in response to Zn intoxication, including the *cyl* gene cluster *cylXDG-acp-cylZABEFIJK* (encoding β-hemolysin/cytolysin) (2.5-4.3-fold down), *lrrG* (25-fold up, leucine rich repeat protein), and *essABC* (11-22-fold down, Type VII secretion system) (Supplementary Table 3). We also note that ~20% of all transcripts (114/567) that were differentially regulated in response to Zn intoxication are predicted to encode hypothetical proteins of unknown function, some of which were up- or down-regulated up to ~29 fold. These observations represent a significant pool of targets for further studies in dissecting bacterial responses to Zn stress.

**Figure 6.**
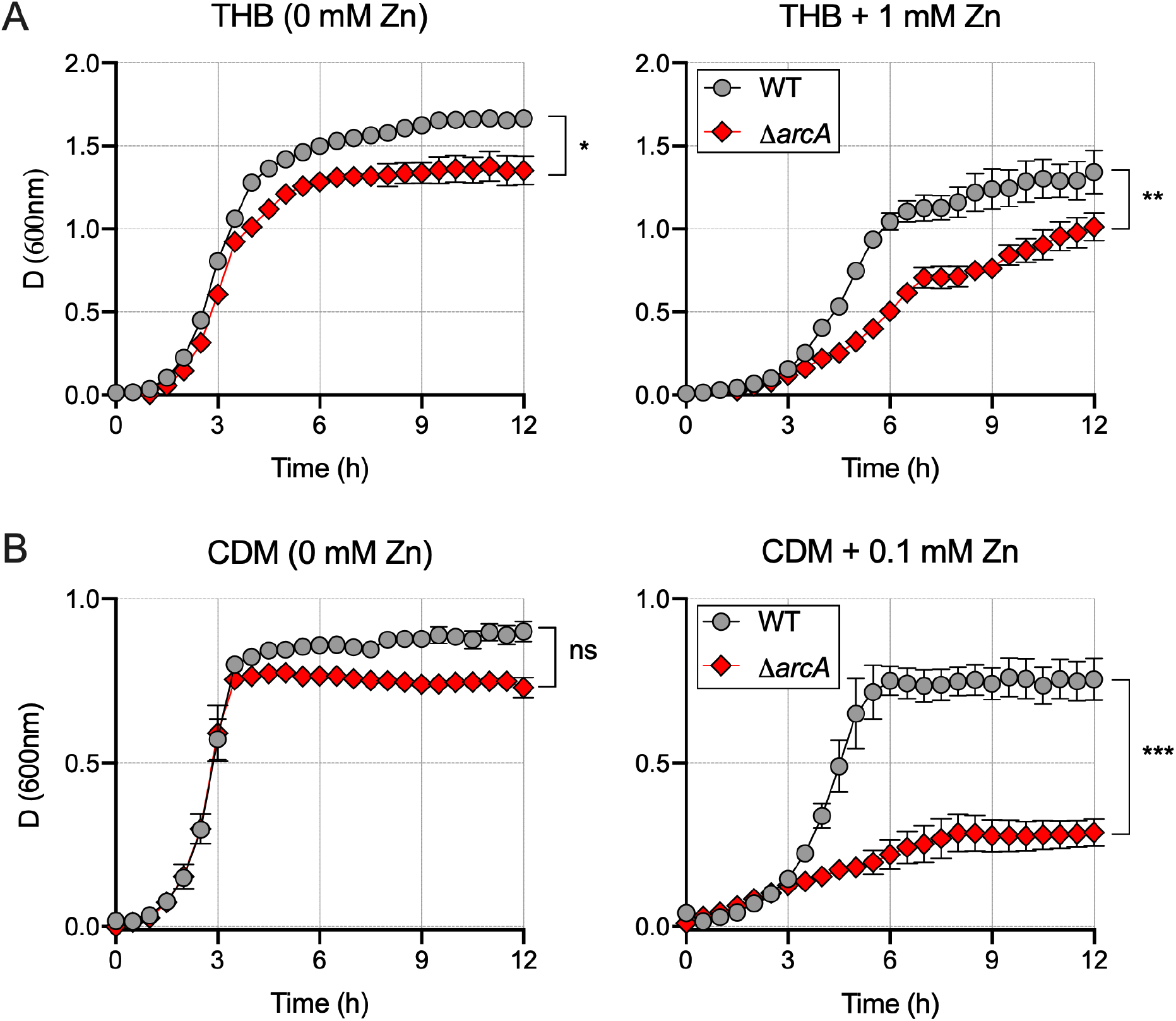
Comparison of the growth WT *S. agalactiae* and an *ΔarcA* isogenic mutant in under Zn stress in A, nutritive THB medium ± 1 mM Zn and B, limiting CDM medium ± 0.1 mM Zn, versus unexposed controls. Data shown are mean measurements of attenuance (*D*; at 600nm) from 3-4 independent experiments, bars show S.E.M and compared using AUC followed by one way ANOVA and Holm-Sidak multiple comparisons (*P < 0.05; **P < 0.01; *** P <0.001).

## Discussion

This study establishes a key role for cellular management of intracellular Zn levels in *S. agalactiae*, via *czcD, sczA* and additional mediators including *arcA*, in conferring an ability to resist Zn intoxication to the bacteria. These findings provide new insight into the molecular mechanisms of virulence used by this pathogen to survive in stressful environmental conditions resulting from elevated metal ion levels, such as within phagocytes during infection in a host. In characterising the Zn efflux systems of *S. agalactiae* in detail, the findings of the present study support prior findings from work on other streptococci (17, 18); for example, our characterisation of regulatory functions of *sczA* that enable cellular management of Zn in *S. agalactiae* are consistent with prior findings reported for *S. pyogenes* (17). These prior observations include that Zn stress upregulates Zn efflux via *czcD* and shuts down Zn uptake via *adcA* (39), with direct effects on the control of intracellular Zn content. Importantly, the Zn stress-response global transcriptome of *S. agalactiae* defined in the current study also elucidates other metal ion transporters and novel additional targets that have not previously been linked to Zn intoxication in any bacteria. A model of the Zn stress response in *S. agalactiae* based on the findings of this study is shown in Figure 7.

**Figure 7.**
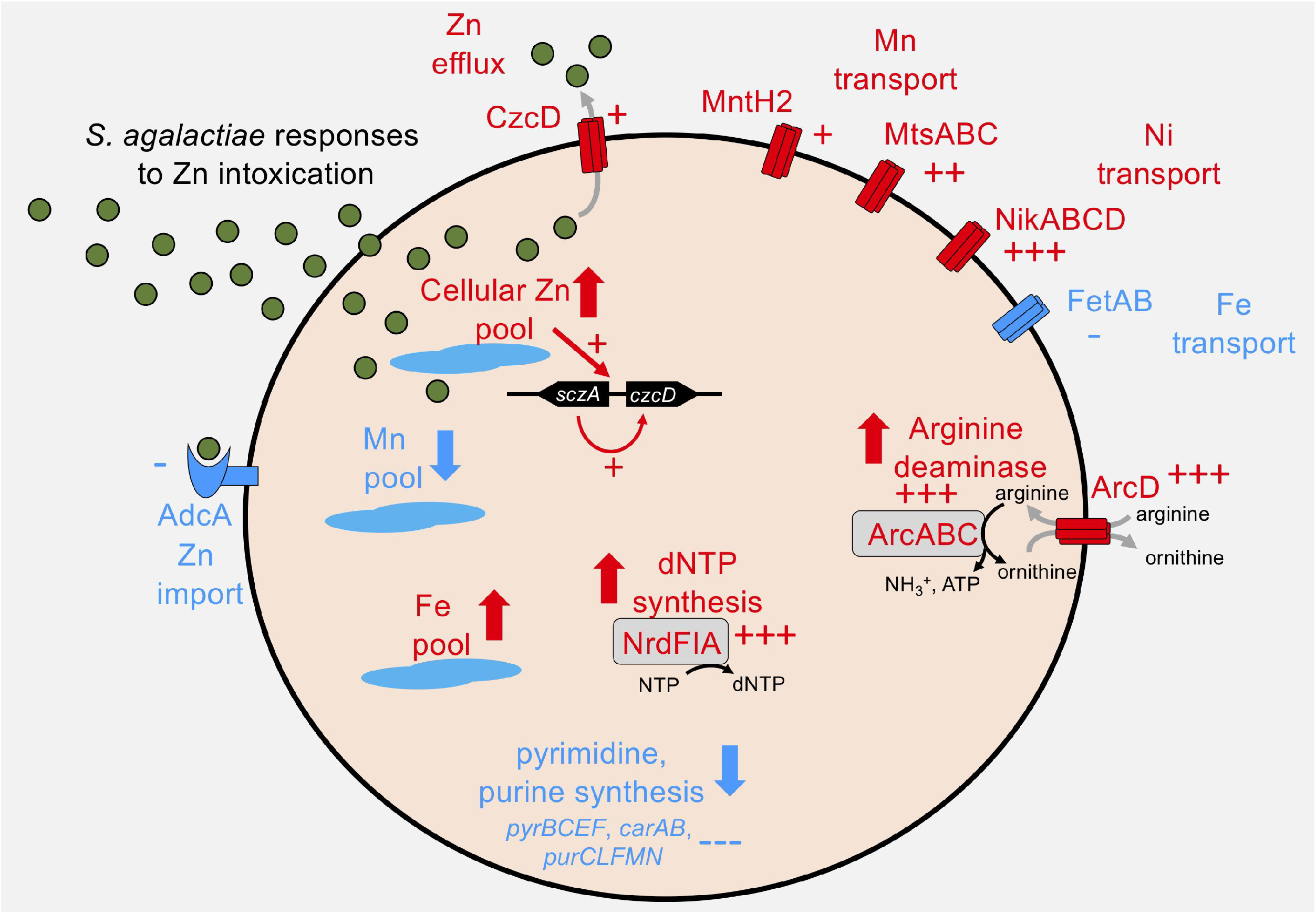
Summary of selected *S. agalactiae* responses to Zn intoxication. *S. agalactiae* senses elevated intracellular Zn to co-ordinate enhanced CzcD expression via the Zn-sensing SczA regulator. Zn stress results in a number of notable changes including differential expression of metal transporters, altered Zn, Mn and Fe cellular pools and metabolic pathways encompassing arginine deaminase, *de novo* purine and pyrimidine syntheses and dNTP synthesis. Red coloring and arrows indicate up-regulation (+ 2-10-fold; ++ 10-20 fold, +++ >20- fold) or down-regulation (- 2-10-fold; --- >20 fold) of a transcript or process.

Systems for Zn acquisition in *S. agalactiae* enable survival of the organism under Zn deficient conditions (9), and confer resistance to calprotectin-mediated metal ion starvation (36) which acts to withhold multiple metals such as Mn and Zn from bacteria in the host (40). Our Zn intoxication transcriptional analyses identify numerous metal transporters that are differentially regulated, including *nikABCD* and *mtsABC*, predicted to transport Ni and Mn, respectively. Given that Ni and Mn import genes have been implicated in bacterial survival during calprotectin stress (36), it is possible that *nik* and *mts* genes respond to altered cellular levels of other metals (rather than Zn) that occur as a consequence of Zn intoxication.

In our study, levels of cellular Mn were perturbed during Zn intoxication. This coincided with corresponding up-regulation of *mntH2* and *mtsABC* transcription, encoding putative Mn transporters. Interestingly, some, but not all *S. agalactiae* strains possess two genes encoding proteins homologous to NRAMP-type MntH transporters; *mntH* (CHF17_00875 = ASZ01148.1) contributes to acid stress responses (41), however a role for *mntH2* (CHF17_02002 = ASZ02237.1) has not been determined. In addition, *S. agalactiae* elevated Fe levels in response to Zn stress, coinciding with down-regulation of a putative Fe export system encoded by *fetA-fetB.* Contrastingly, Cu levels and expression of *copA* (encoding a putative Cu transporter) were unaffected in *S. agalactiae* exposed to Zn stress. Collectively, these data show broader dysregulation of metal management in *S. agalactiae* undergoing Zn intoxication beyond Zn itself, encompassing Mn and Fe, consistent with a prior report in pneumococci (14). Future examination of the roles of the Ni, Mn, Fe and Cu transport in *S. agalactiae* in the setting of bacterial metal stress and in models of infection would therefore be of interest.

The finding that numerous core metabolic pathways in *S. agalactiae* are impacted by Zn stress, including some related to synthesis of purines, pyrimidines, riboflavin and dNTPs, underscores the critical nature of Zn management for maintenance of basic cellular processes in the bacteria. The arginine deaminase system was among the most differentially regulated gene clusters, undergoing 20-30-fold enhanced expression during Zn stress. Remarkably, *arcA*-deficient *S. agalactiae* exhibited heightened sensitivity to Zn stress, establishing a novel role for this system in streptococcal resistance to metal ion intoxication. The precise molecular mechanisms that underpin how this system renders the bacterial cell resistant to Zn intoxication requires further definition but arginine deaminase typically converts arginine to ammonia, ATP and ornithine; overproduction of one of these may support bacterial survival during Zn stress. A recent study identified arginine deaminase as a key factor in resistance to antibiotics and biofilm formation in *S. pyogenes* (42). Numerous prior studies have reported elevated expression of *arc* genes in *S. agalactiae* in response to acid stress (43), human serum (44), blood (45), or amniotic fluid (46). This suggests important but yet to be defined roles for arginine deaminase in virulence of *S. agalactiae.* Together with the findings of the current study, these observations suggest that arginine deaminase supports the ability of *S. agalactiae* to respond to diverse stressors in addition to Zn intoxication.

Analysis of the role for Zn efflux systems of *S. agalactiae* in pathogenesis showed no contribution to bacterial survival in macrophages. This finding is surprising in the context of a prior study of *S. pneumoniae sczA* (that is functionally analogous to *S. agalactiae sczA)*, which reported a major role for Zn cellular management via *sczA* in the intracellular survival of the bacteria in human macrophages (35). We confirmed the generation of a robust Zn mobilization response in the host cells following *S. agalactiae* infection but observed equivalent fitness of mutants for Zn management compared to WT for survival in murine and human macrophages. Considering the conserved nature of *szcA* between streptococci, it is plausible that these mutants might be attenuated in other phagocytes such as neutrophils, as described for *S. pyogenes* (17), noting a need for further examination of differences in metal ion resistance phenotypes among streptococcal species.

The discovery of attenuation for colonization in a mouse model of disseminated infection in the *S. agalactiae* mutants in Zn management systems in this study establishes a role for Zn stress in disease pathogenesis due to this organism. The attenuations observed for the *czcD^-^* and *sczA^-^* mutants in the liver and bladder, and the blood and heart, respectively, were not dramatic according to the tissue bacterial loads subsequent to blood challenge; however, the attenuations were statistically significant; and the phenotypes observed were also specific to these tissues because multiple other tissue-types that were analysed had equivalent numbers of WT bacteria and mutants. Considered together, these findings point to tissue-specific effects of the ability to manage Zn stress in *S. agalactiae* colonization of the mammalian host. Prior studies have shown more dramatic effects of *czcD* and *sczA* mutations in reducing bacterial virulence in mouse models of infection. For example, *czcD* mutation rendered *S. pyogenes* non-lethal in a subcutaneous infection model in mouse (17). *czcD* mutation attenuated *S. pneumoniae* 2.6-fold for survival in the lungs of mice following intranasal challenge compared to WT bacteria (47). Evaluating the current findings in the context of these prior studies highlights the differences in experimental designs viz. types of infection models, bacterial species and outcome measures that would likely influence any potential degree of attenuation in mutants (17, 35, 47). More broadly, this highlights the importance of evaluating the roles of individual mediators such as CzcD, SczA and ArcA in appropriate experimental models that are designed to parallel natural infection in the human host. For *S. agalactiae*, for example, exploration of the role of resistance to Zn stress in colonization of the female genital tract (48), and the brain (49) would be of interest given the propensity of this pathogen to cause these infections in humans (50).

In conclusion, this study identifies new mediators of Zn cellular management in *S. agalactiae*, and shows that resistance to Zn stress in this pathogen contributes to colonization in the host. Future examination of these mediators, and their role survival of *S. agalactiae* and other bacterial pathogens in relevant settings of infection will be important to more fully understand microbial resistance to metal intoxication and it influence on virulence and pathogenesis.

## Acknowledgments

We thank Michael Crowley and David Crossman of the Heflin Centre for Genomic Science Core Laboratories, University of Alabama at Birmingham (Birmingham, AL) for RNA sequencing. We thank Timothy Barnett who provided pLZ12 containing cloned mCherry. We also thank Lahiru Katupitiya and Dean Gosling for excellent technical assistance and Ryan Stewart at the School of Environment Analytical Chemistry Core Facility, Griffith University, for ICP-OES. This work was supported by a Project Grant from the National Health and Medical Research Council (NHMRC) Australia (APP1146820 to GCU).

